# Triplet-based similarity score for fully multi-labeled trees with poly-occurring labels

**DOI:** 10.1101/2020.04.14.040550

**Authors:** Simone Ciccolella, Giulia Bernardini, Luca Denti, Paola Bonizzoni, Marco Previtali, Gianluca Della Vedova

**Affiliations:** Department of Computer Systems and Communication, University of Milano-Bicocca, Milan, Italy

## Abstract

The latest advances in cancer sequencing, and the availability of a wide range of methods to infer the evolutionary history of tumors, have made it important to evaluate, reconcile and cluster different tumor phylogenies.

Recently, several notions of distance or similarities have been proposed in the literature, but none of them has emerged as the golden standard. Moreover, none of the known similarity measures is able to manage mutations occurring multiple times in the tree, a circumstance often occurring in real cases.

To overcome these limitations, in this paper we propose MP3, the first similarity measure for tumor phylogenies able to effectively manage cases where multiple mutations can occur at the same time and mutations can occur multiple times. Moreover, a comparison of MP3 with other measures shows that it is able to classify correctly similar and dissimilar trees, both on simulated and on real data.

## 1 Introduction

Recent methods to accurately infer the clonal evolution and progression of cancer have made it possible to develop targeted therapies for treating the disease. As discussed in several studies [1, 2], understanding the history of accumulation and the prevalence of somatic mutations during cancer progression is a fundamental step to devise these treatment strategies.

Given the importance of the task, a multitude of methods for cancer phylogeny reconstruction have been developed over the years. The increasing number of tools created has been encouraged by the diversity of data available; for instance, we are witnessing a shift from bulk sequencing data [3, 4, 5, 6, 7] towards single-cell data [8, 9, 10, 11] and hybrid approaches [12, 13].

Having many different tools accomplishing the same task requires solid methods to compare their results. In contrast with classical phylogenetic trees, whose leaves, and only leaves, are labeled (with the species they represent), the trees that model tumor phylogenies are *fully-labeled*, i.e., they also have labels (corresponding to the mutations) on the internal nodes. While there is a wide range of measures to compare leaf-labeled trees in the literature, ad-hoc methods for tumor phylogenies are starting to appear in the last few years [14, 15, 16, 17, 18]; in particular, a detailed study of some notions of distance [14] has introduced two new measures complementing some more established definitions used in various cancer inference studies [19, 9]. Those new measures are more nuanced, in order to capture some aspects of the mutation inheritance process, while still being very efficient to compute. A common trait of all the latter distances is their reliance on the analysis of *pairs* of nodes.

On the other hand, some of the most widely used distances on classical phylogenies are based on rooted triples [20, 21, 22] (for rooted phylogenies) or quartets [23] (for unrooted phylogenies) of labeled leaves. Although such metrics have major limitations for our purposes, as they do not apply directly to fully-labeled trees, they also have some desirable properties that we would like to transfer in our setting. Specifically, this kind of metric captures well the differences in the topology of the trees; a feature that, to the best of our knowledge, lacks in most of the existing methods for tumor phylogenies. Therefore we expect a triplet-based measures to provide additional insights on the different evolutionary histories, when applied to cancer progression.

In this paper, we generalize the notion of rooted triples similarity for classical phylogenies to tumor phylogenies. Moreover, we further extend this to multi-labeled trees (that is, where each node is labeled by a set of labels) and poly-occurring labels (that is, each label can be assigned to more than one node). The latter feature is needed since recent studies [24, 25] suggest widespread recurrence and loss of mutations, and more and more methods designed to infer tumor phylogenies considering such a possibility are starting to appear [11, 19, 9]. In a phylogenetic tree a mutation loss is represented by a special character in the label, such as a minus sign: the design of our measure allows to handle such evolutionary events effectively, as they uniquely correspond to their label like any other kind of mutation.

Through an extensive experimental analysis, we show that our novel measure is able to overcome the limitations in the existing literature and to provide a better alternative to both the direct comparison of evolutionary histories and the application to established clustering techniques, following the approach of [14]. Such a performing measure can also be incorporated in recent works [26, 27] designed to cluster and build consensus across multiple cancer progressions. An open source implementation of MP3 is publicly available at https://github.com/AlgoLab/mp3treesim.

## 2 Methods

A classical phylogenetic tree is a rooted, unordered, leaf-labeled tree. The set of all the labels occurring in *T* is denoted by *λ*(*T*), and a function N(·) maps each element of *λ*(*T*) to a leaf of *T*. We denote with `LCA`(*u, v*) the Lowest Common Ancestor of nodes *u* and *v*. Given three leaves *u, v, z* ∈ *V*_*T*_, the *minimal tree topology* they induce on *T*, denoted as `MTT`_*T*_ (*u, v, z*), is the smallest subtree of *T* that includes the nodes 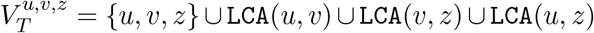, and where all the nodes with degree 2 not in 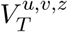 are contracted.

The rooted triplet distance measures the dissimilarity between two leaf-labeled trees with identical labels. It is given by the number of rooted triplets that induce different minimal topologies (Figure 1) in the two trees over the total number of triplets [28]. As tumor progression trees are fully-labeled, such metric cannot be directly applied: in this section we propose a novel similarity measure, inspired by the triplet distance, specifically designed for these more general trees.

**Figure 1:**
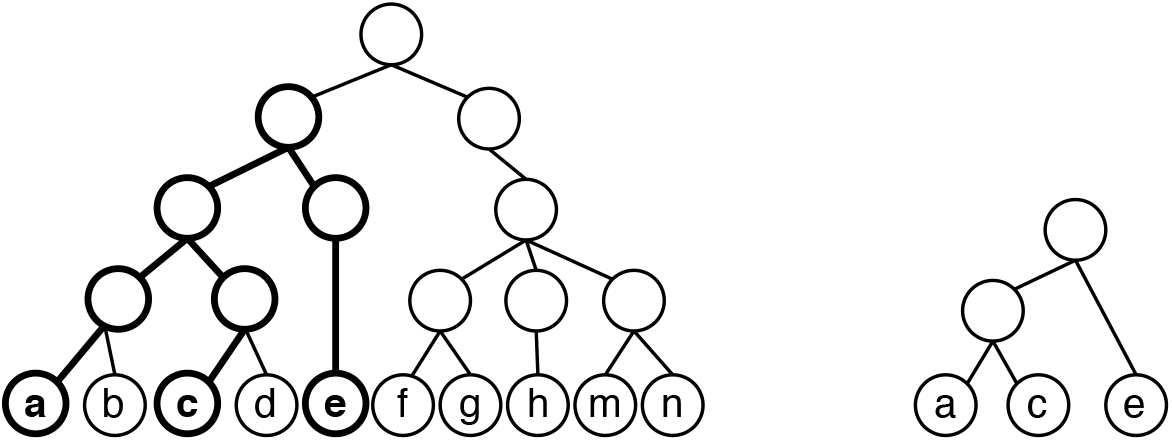
Rooted triplet on labels (*a, c, e*). (*Left*) Tree *T* where the smallest subtree that contains all three labels is highlighted. (*Right*) The minimal topology induced by (*a, c, e*).

### 2.1 Extension to fully labeled trees and multi-labeled trees

A tree *T* on a set *V*_*T*_ of *n* nodes is *fully-labeled* by a set *λ*(*T*) of labels if there is a bijection N : *λ*(*T*) → *V*_*T*_. The definition of minimal topology of three leaves can be trivially extended to the minimal topology of three nodes: we next show that there are only five possible configurations (see Figure 2).

**Figure 2:**
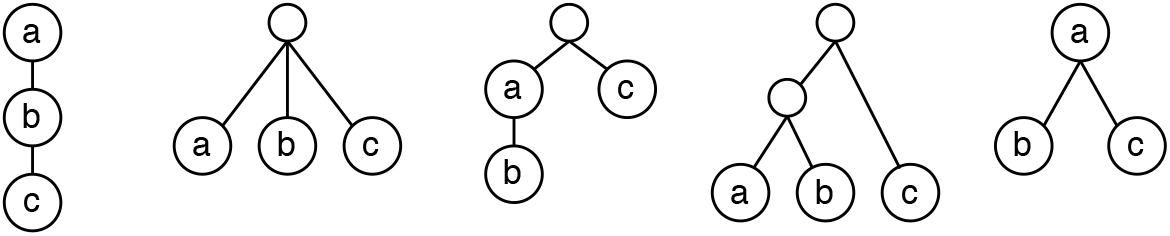
The five possible configurations for the minimal tree topology induced by three nodes.

#### Lemma 1.

*Given nodes u, v, z* ∈ *V*_*T*_, *there exist only five possible configurations for* `MTT`_*T*_ (*u, v, z*).

*Proof.* We start by dividing two possible cases: (i) `LCA`(*u, v*) = `LCA`(*v, z*) = `LCA`(*u, z*), or (ii) just two `LCA`s are the same, say `LCA`(*v, z*) = `LCA`(*u, z*). There are no other possibilities, as `LCA`(*u, v*) ≠ `LCA`(*v, z*) ≠ `LCA`(*u, z*) is impossible: indeed, suppose without loss of generality that `LCA`(*u, v*) is a descendant of `LCA`(*u, z*), `LCA`(*u, v*) ≠ `LCA`(*u, z*): they cannot be unrelated, as by definition they are both ancestors of *u*. `LCA`(*u, z*) is thus a common ancestor for *v* and *z*. Suppose towards a contradiction that `LCA`(*v, z*) ≠ `LCA`(*u, z*), thus it is a descendant of `LCA`(*u, z*) and an ancestor of `LCA`(*u, v*). But then it is an ancestor of both *u* an *z* and it is lower than `LCA`(*u, z*), a contradiction.

Case (i) has two subcases: either `LCA`(*u, v*) ∈ {*u, v, z*}, corresponding to the rightmost configuration in Figure 2, or `LCA`(*u, v*) ∈*/* {*u, v, z*}, corresponding to the second configuration from the left. Case (ii) has three subcases: either both the distinct `LCA`s are in {*u, v, z*}, or none of the two is, or finally one is in {*u, v, z*} and the other is not. The first subcase corresponds to the leftmost configuration in Figure 2, the second subcase to the fourth configuration from the left. For the third subcase, either the external `LCA` is an ancestor of all of the three {*u, v, z*}, corresponding to the third configuration, or it is an ancestor of two nodes and a descendant of the third one, say *u*. In the latter case, though, the external node would be the only child of *u*, and thus would be contracted by definition of `MTT`_*T*_ (*u, v, z*), leading again to the rightmost configuration of Figure 2.

In the case of fully-labeled trees, the definition of `LCA` of two nodes and `MTT` of three nodes can trivially be extended to the `LCA` of two labels and the `MTT` of three labels, as there is a one-to-one correspondence between nodes and labels. From now on, for ease of presentation, given two nodes *u* and *v* and their respective labels *a* and *b*, we will use `LCA`(*u, v*) and `LCA`(*a, b*) interchangeably. When modeling tumor progression, though, to have a bijection between nodes and labels (i.e., mutations) is quite a strong assumption, as multiple mutations often appear at the same time in the evolutionary history of cancer. We thus relax our assumptions and consider *multi-labeled* instead of fully-labeled trees.

A rooted, unordered tree *T* is multi-labeled if there exists a surjective function N : *λ*(*T*) → *V*_*T*_ that labels each node of *T* with a set of labels from *λ*(*T*): note that, in this model, each label is assigned to one and only one node of *T*. We extend the definition of lowest common ancestor of two labels for a multi-labeled tree as follows: if *a* ∈ *λ*(*T*) and *b* ∈ *λ*(*T*) label the same node *u*, then `LCA`(*a, b*) = *u*; if they label two distinct nodes *u, v*, then `LCA`(*a, b*) = `LCA`(*u, v*). This allows us to straightforwardly extend the definition of minimal tree topology of three labels for multilabeled trees. There are only four possible additional configurations for the minimal tree topology of multi-labeled trees, shown in Figure 3: a proof can be found in the Supplementary Materials.

**Figure 3:**
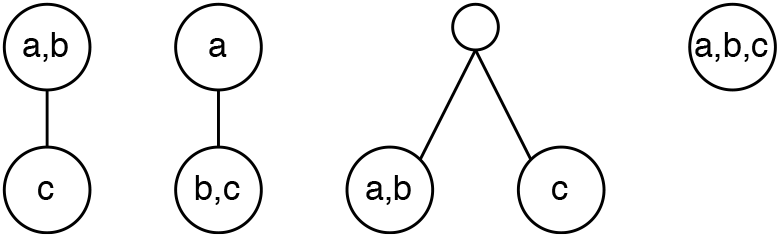
The four additional possible configurations for the minimal tree topology of multi-labeled trees induced by three nodes.

#### Lemma 2.

*Given T multi-labeled and a, b, c* ∈ *λ*(*T*), *there exist nine configurations for* `MTT`_*T*_ (*a, b, c*).

### 2.2 Extension to poly-occurring labels

We further extend our model of tumor phylogeny by allowing the same label of *λ*(*T*) to be assigned to multiple nodes of *T*. An element of *λ*(*T*) that labels more than one node of *T* is said to be a *poly-occurring* label. To the best of our knowledge, none of the existing tools is able to handle poly-occurring labels: indeed, although some of them accept input trees with poly-occurring labels, they simply disregard the multiple occurrences of a same label.

Since it is often the case where the inferred evolutionary history involves the appearance of the same mutation in multiple events, a meaningful comparison between tumor phylogenies cannot overlook such a phenomenon. To consider poly-occurring labels in our similarity measure, we extend the definition of minimal tree topology. First, note that if a label occur multiple times in the tree, then N maps each label to one or more nodes in *V*_*T*_. Then, we define the minimal tree topology of poly-occurring labels *a, b, c*, denoted by `M`, as follows, where ⊔ indicates the multiset union:

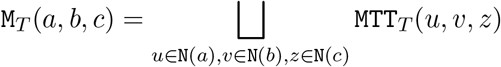

In other words, the minimal tree topology of three labels is the multiset of all the minimal tree topologies of the nodes where *a*, *b*, and *c* appear. We remark that in this setting `M`_*T*_ is a multiset of configurations, thus the same configuration may appear multiple times in `M`_*T*_.

### 2.3 Similarity measure between trees

We are now able to define a similarity measure between fully-labeled trees with poly-occurring labels. Let *S* be a multiset and let |*S*| be its cardinality. We define the number of shared configurations of labels *a, b, c* between two trees *T*_1_ and *T*_2_ as 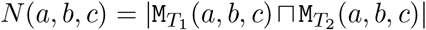, *i.e.* the cardinality of the multiset intersection, and the maximum number of configurations of the triplet in the trees as 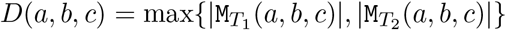.

Based on these two values we define multiple variations of the Multi Poly-occurring labels triplet-based (`MP3`) similarity measure that we will later combine into a single score. We define `MP3`_*⋂*_ as the similarity computed between triplets of labels shared by the two trees:

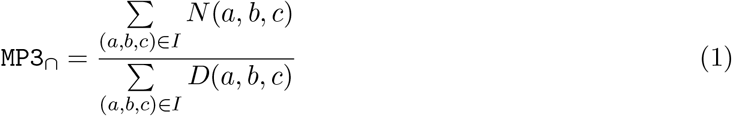

where *I* is the set of triples in *λ*(*T*_1_) ∩ *λ*(*T*_2_). Due to the nature of only considering the subset of labels that appears in both trees, `MP3`_*⋂*_ is a conservative measure, therefore we present a variation that consider all possible configurations in both trees, thus having a wider view:

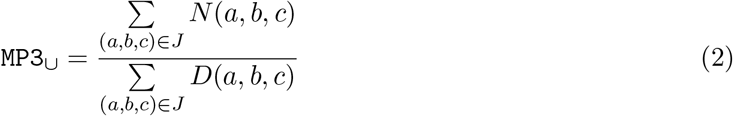

where *J* is the set of triples in *λ*(*T*_1_) ⋃ *λ*(*T*_2_). Differently from `MP3`_*⋂*_, `MP3`_*⋃*_ weighs also the the labels that appear only in one of the trees. Note that, for every pair of trees, `MP3`_*⋃*_ ≤ `MP3`_*⋂*_, as the numerator remains identical in both, while the denominator of `MP3`_*⋃*_ has all the elements in `MP3`_*⋂*_ with the addition of the values of *D* for the triples present only in one of the input trees.

Although `MP3`_*⋂*_ and `MP3`_*⋃*_ are closely related, they provide two different views of a tumor phylogeny. Indeed, on one hand `MP3`_*⋂*_ measures how similar the shared history of two tumor phylogenies is, *i.e.* it provides an idea of how well the two progressions can be reduced to the same subsequence of common mutations. On the other hand, `MP3`_*⋃*_ measures how similar the whole history of the two evolutions are, *i.e.* it considers the impact of mutations acquired only in one progression.

Since the previous measures capture different aspects of the progressions, we want to combine them into a single, usable and powerful similarity measure that couples the strengths of both. The most intuitive method is to simply use a mean. We opted for the geometric mean: 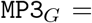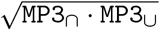

This function is not completely satisfactory, as a uniform function of 1 and 2 is not able to comprehensively capture the nuances in the input trees. Therefore we developed a weighted mean with an intentional bias towards `MP3`_*⋂*_ to catch inner similarities in different trees. Such combination then tends to be closer to `MP3`_*⋂*_ when the trees are similar while moving towards `MP3`_*⋃*_ as the trees are less similar:

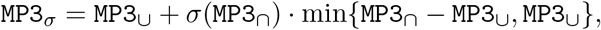

where 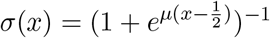 is the classic sigmoid function centered in 1/2 and *μ* is used to adjust the slopeness of the curve; we set *μ* = 10 in our experimentation. In addition the sigmoid polarizes the values close to 1/2, thus helping decide whether they are closer to 1 or 0, therefore moving the final score closer to `MP3`_*⋂*_ or `MP3`_*⋃*_.

While all four measures are available in our implementation, we decided to use `MP3`_*σ*_ as default measure and is denoted simply as `MP3`. An experimental comparison of all four measures is shown in the Supplementary Materials.

## 3 Results

### 3.1 Simulated Data

To perform our experiments we follow an approach similar to the one performed in [14]. We start from a base tree on which we apply a series of perturbations selected from: label swapping, label removal, label duplication, node swapping and node removal. Both the perturbations and the nodes and labels on which they are applied are chosen at random: our procedure allows to select a user-specified total number of actions and a probability vector that will be used to select the perturbations from the previous list.

For the measure comparison experiments, we generated 30 perturbations from each of the 5 base trees, for a total of 150 trees. For the clustering evaluation, 3 base trees are entirely different from each other, and another 2 are perturbations of two of the others, to simulate similar sub-families of the same tumor type: we perform a total of 10 perturbations on such 5 trees. More details on the perturbation parameters will be described in each section, while the entire configuration is available and reproducible at https://github.com/AlgoLab/MP3treesim_supp.

### 3.2 Measures comparison

We compared `MP3` against all the different versions of DISC and CASet from [14] and MLTD [15]. While `MP3` and MLTD provide similarity scores, DISC and CASet compute a dissimilarity score, that we convert into a similarity measure by simply subtracting their value from 1.

#### 3.2.1 Effect of changes in the tree topology

A key feature a measure on tumor phylogenies should have is to DISCern changes at different tree depths; indeed, a change close to the root should be more impactful than a change towards the leaves. Such a behavior is fundamental, as driver mutations are often acquired early in the evolutionary history, while less important passenger mutations usually happen at later stages: to mistake the two types of mutations should therefore have a high impact on a good similarity measure.

To estimate this effect on all the measures, we start from a linear base tree (T_0_ in Figure 4 (*Left*)); we then raise its only leaf one level at the time and compute its similarity to the base tree, expecting a drop in similarity as the leaf raises to the root, similarly to experiment proposed in [14]. Figure 4 (*Left*) clearly displays such effect for `MP3`, showing that it has the highest similarity decrease among all measures; DISC and CASet also have similar trends, but to a lower extent. Since the set of labels is the same for all trees, there is no difference between union and intersection versions of DISC and CASet. Contrarily, as already observed in [14], MLTD plateaus after the first change.

**Figure 4:**
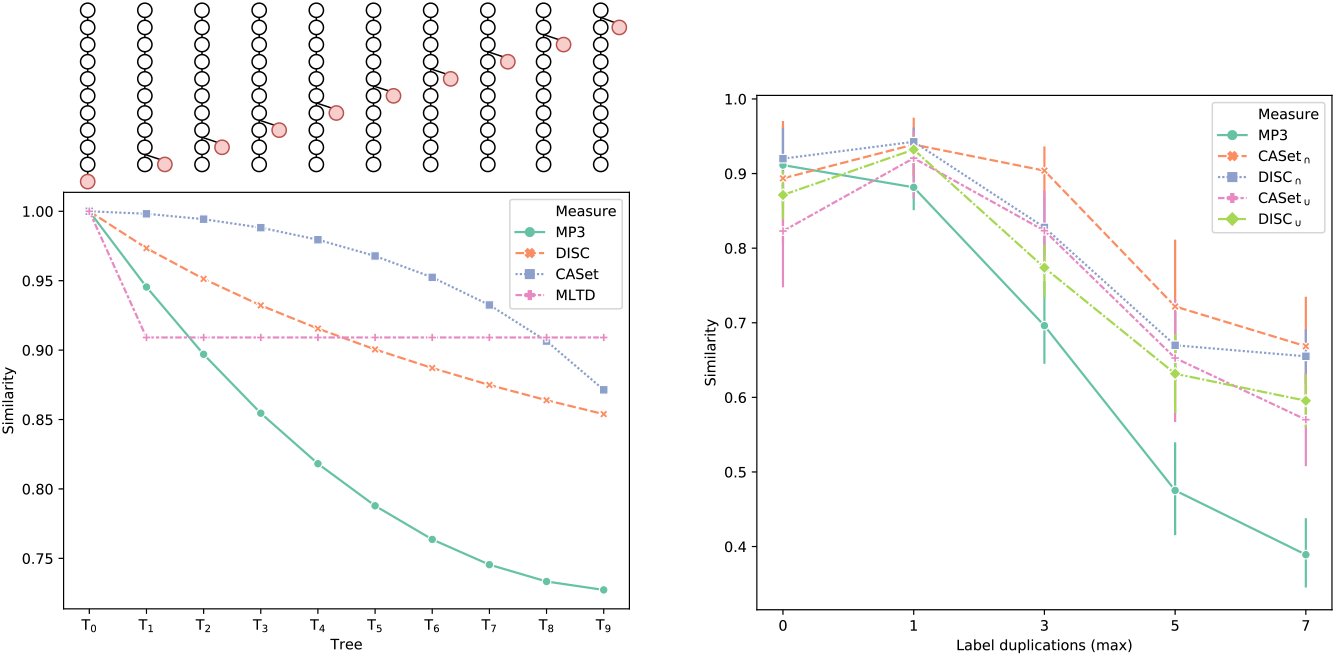
(*Left*) Effect of a node (highlighted in red) that ascends from leaf to child of the root, T_0_ is the base tree to which the others are compared. (*Right*) Effect of label duplication on the similarity scores. Similarities are the average of 15 trees generated from the same base with the specified maximum number of duplications. MLTD was excluded since it failed to run on instances with poly-occurring labels.

Another interesting aspect to investigate is how the presence of poly-occurring labels influences the similarity scores, as the more sophisticated the inference tools get, the more is common to have tumor phylogenies with multiple acquisitions or losses of the same mutation. To evaluate this aspect we started from a multi-labeled base tree with all labels occurring only once. We then created 15 perturbed trees for 5 different configurations. In the first one (on the abscissa 0 in Figure 4 (*Right*)) we allowed one operation excluding label duplication; for the others we allowed a total of 1, 3, 5 and 7 operations with much higher chance of selecting a label duplication. Since perturbations occur randomly, we are only sure that at most the specified number of duplication occurred, and not necessarily to the same label.

Figure 4 (*Right*) shows that CASet_*⋂*_, CASet_*⋃*_, DISC_*⋂*_ and DISC_*⋃*_ have similar trends in this setting, `MP3` being the only one that differs. In particular, the other measures assign an higher similarity score to the second configuration than to the first one, despite they are both obtained with one perturbing operation, allowing label duplication only in the second one. `MP3` is the only measure that positively displays a monotonic decrease in similarity as the number of poly-occurring labels increases, being markedly steeper than the others. We believe that a larger steepness will be more informative than a plateauing curve, since while being true that after many of poly-occurrences no more information is gained, all the duplications will inevitably add more and more noise to the tree. Since MLTD assumes that every label appears only once, it failed to run on this experiment and was therefore excluded.

#### 3.2.2 Results on simulated data

To analyze the differences between all measures we designed two experimental settings: from 5 different base trees (available in the Supplementary Materials) we generated 30 perturbations for each class and computed similarities scores between all the 150 resulting trees. In the first configuration we allowed a total of 3 operations excluding label duplications, while in the second one we allowed them. All the parameters and the different probabilities used for applying perturbations are available at our supplementary repository https://github.com/AlgoLab/MP3treesim_supp.

Results for the first configuration are shown in Figure 5. The heatmaps (*Left*) show that `MP3` discerns the best between the trees in the same class (main diagonal) and the others: the results of DISC_*⋃*_ are really close to ours, but there is a more noticeable noise outside the main diagonal. DISC_*⋂*_ and CASet_*⋃*_ present even more noise than the others, but are still mostly able to distinguish the different classes; CASet_*⋂*_ seems to struggle the most on this setting, while MLTD displays high values of similarities for every couple of trees, but it is still able to differentiate between the bases.

**Figure 5:**
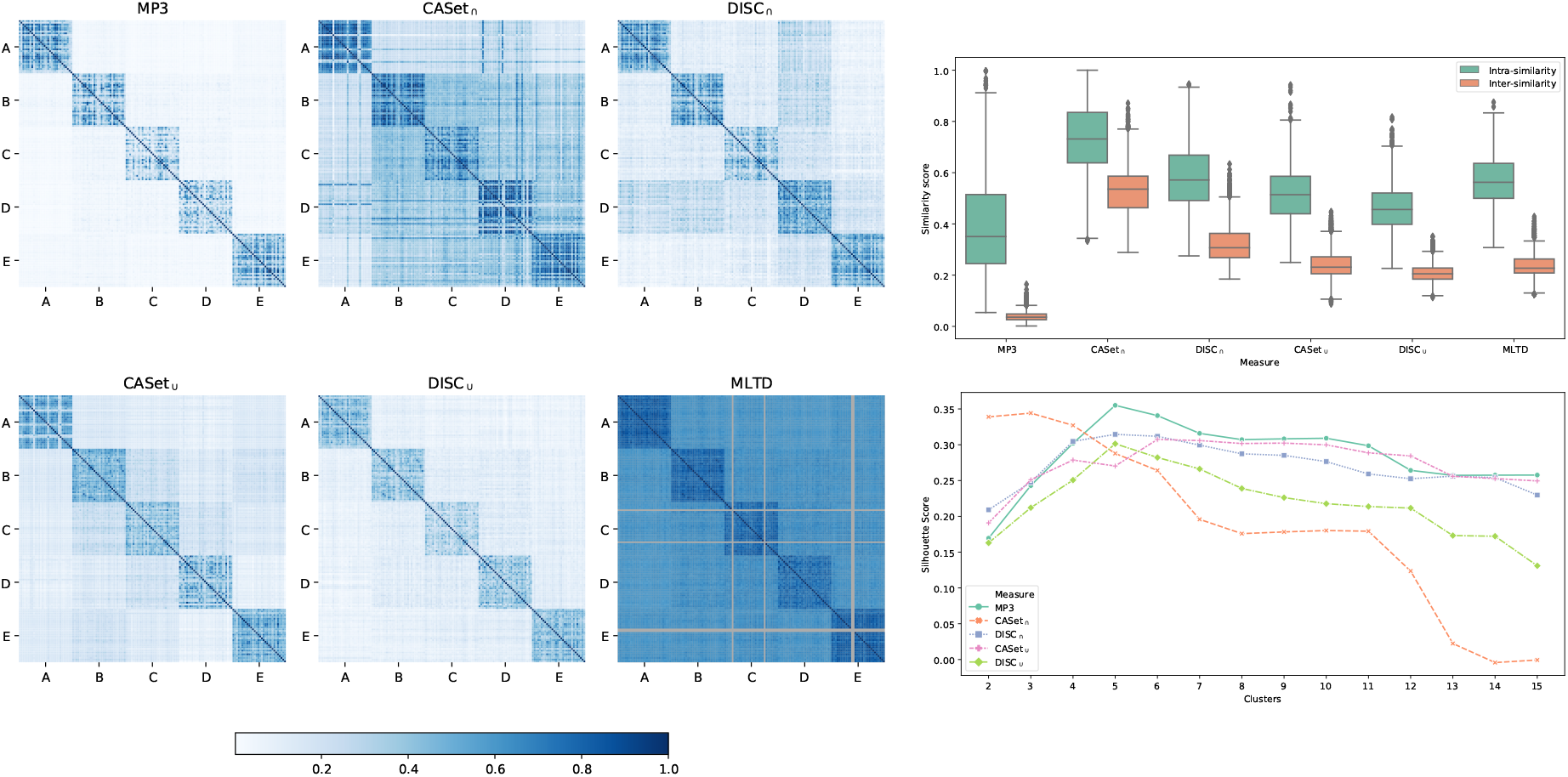
Results for the first experimental configuration: (*Left*) Heatmaps displaying the scores between all-pairs 150 simulate trees. (*Top-Right*) Distribution of the similarities between the trees in the same class (Intra-similarity) and in different classes (Inter-similarity). (*Bottom-Right*) Silhouette score computed using a hierarchical linkage clustering with cuts from 2 to 15.

The boxplots in Figure 5 (*Top-Right*) show the same result quantitatively: the crucial feature is to correctly distinguish the different classes. The values represent the distribution of the similarities between the trees in the same class (Intra-similarity) and in different classes (Inter-similarity). `MP3` differentiates better between intra and inter similarity, exhibiting the most compact distribution for the inter-similarities scores, while being a little more dispersed on the intra-similarity due to the action of the sigmoid, that pulls apart the values around 1/2. Similarly to the previous case, DISC_*⋃*_, CASet_*⋃*_ and MLTD show similar trends, while CASet_*⋂*_ displays the most overlapping distributions.

Lastly, in Figure 5 (*Bottom-Right*), we computed a silhouette score from the data using a hierarchical linkage clustering with cuts from 2 to 15 to simulate a clustering scenario. Once again, `MP3` performs the best expressing the maximum value for 5 cuts, being the 5 classes. DISC_*⋂*_, DISC_*⋃*_ also show the largest value at the same cut. MLTD was excluded from the plot since it scored values close to −1 for every cut, thus causing the figure to be hard to interpret.

In the second experimental setting we introduced poly-occurring labels to the simulation. Figure 6 exhibits results very similar to the previous ones. The main difference is that in the silhouette score (*Bottom-Right*) `MP3`, while still having its maximum value in correspondence of 5 cuts, is slightly lower than the other measures. On this experiment MLTD, not allowing poly-occurring labels, failed to compute the score in most of the instances, shown in grey in the heatmaps (*Left*); it was excluded from the other plots given the high amount of failed runs.

**Figure 6:**
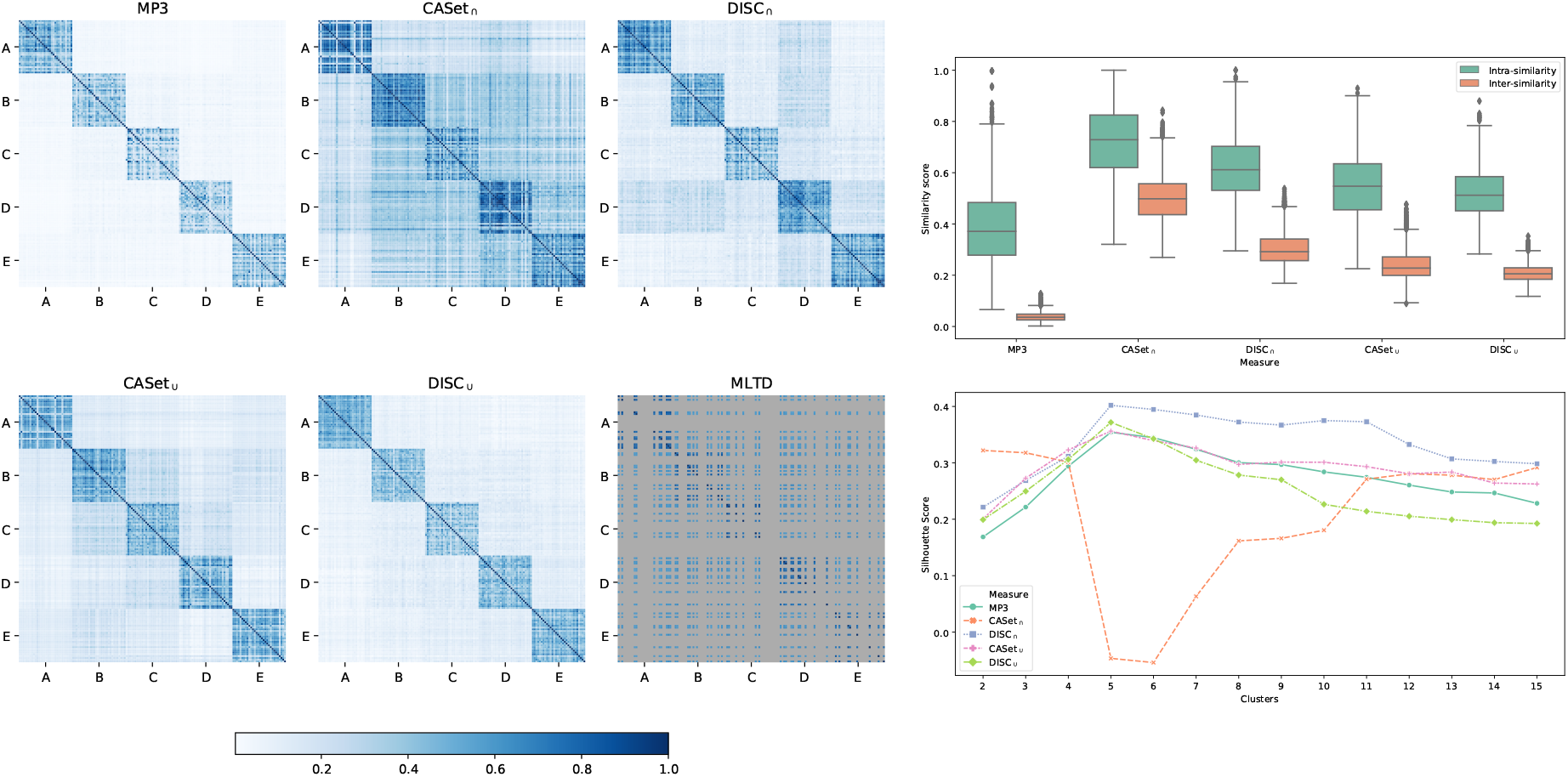
Results for the second experimental configuration: (*Left*) Heatmaps displaying the scores between all the 150 simulate trees. (*Top-Right*) Distribution of the similarities between the trees in the same class (Intra-similarity) and in different classes (Inter-similarity). (*Bottom-Right*) Silhouette score computed using a hierarchical linkage clustering with cuts from 2 to 15.

### 3.3 Application to clustering of trees

A very important application of a tree similarity measure is clustering, e.g., to classify cancer type of patients by the similarity of their inferred phylogenies. This is of crucial interest for the development of precision therapies based on the topological structure and the evolution of mutations. Since to curate such classifications manually would be unfeasible as the size and the number of mutations increases, a good measure to use in conjunction with a clustering method is necessary.

To evaluate a similar scenario we started from 3 different bases, then perturbing two of such trees chosen at random; these new trees are then considered as additional base trees. Given this 5 bases we created a total of 10 perturbed trees from each class. The goal was to simulate an experiment with three separate classes, with two of them further split in two subclasses, to obtain subtypes of the same cancer families. The five resulting bases are available in the Supplementary Materials and the parameters used for the simulations are in our supplementary repository.

Results for the clustering experiment are reported in Figure 7; (*a*) shows the clustermaps computed using hierarchical linkage clustering. `MP3`, DISC_*⋂*_ and DISC_*⋃*_ correctly cluster the three main families as well as the two sub-families, while both versions of CASet struggle the most in this experiment. Figure 7 (*b*) displays the distribution of intra- and inter-similarity between the five bases; `MP3` has the most compact inter-similarity distribution and is the only method that completely separates intra- and inter-distributions. The high number of outliers for all methods is due to the high similarity of the two subclasses. To confirm this hypothesis we computed the same distributions only for the three main classes, remapping the subclasses to the original corresponding base class in (*d*), where we note that the number of outliers is significantly reduced. Finally, Figure 7 (*c*) shows the silhouette scores for the dataset; all measures express a higher score with 3 cuts, suggesting that the two subclasses are very similar to the two main bases they are derived from. The scores are very similar for all measures, with DISC_*⋃*_ having a higher value with 3 cuts and `MP3` having a slightly higher with 5 clusters. CASet_*⋂*_ is the only method that have a much higher score in 5, however, as shown in (*a*), the five clusters it reports are not the correct ones. MLTD was excluded from this experiment because it failed to run on most instances due to poly-occurring labels.

**Figure 7:**
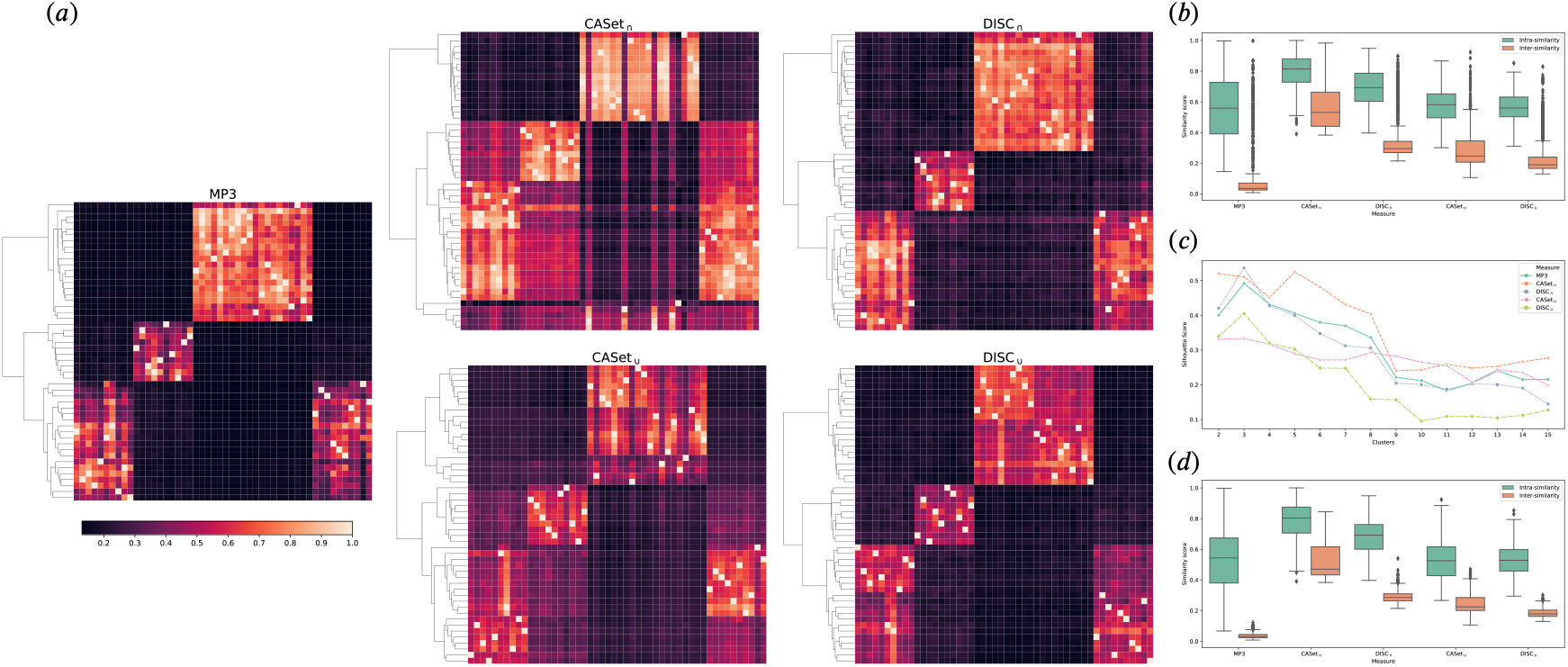
Results for the clustering experiment: (*a*) Clustermaps of the 50 simulated trees computed using hierarchical linkage clustering. (*b*) Distribution of the similarities between the trees in the same class (Intra-similarity) and in different classes (Inter-similarity) for the 5 classes. The high number of outliers for all methods is due to the high similarity of the two subclasses. (*c*) Silhouette score computed using a hierarchical linkage clustering with cuts from 2 to 15. (*d*) Distribution of the similarities between the trees in the same class (Intra-similarity) and in different classes (Inter-similarity) for the three main classes, remapping the subclasses to the original corresponding base. MLTD was excluded from this experiment because it failed to run on most instances due to the presence of poly-occurring labels.

### 3.4 Application to real dataset

To further evaluate our similarity measure, we applied it to two publicly available real datasets: breast cancer xenoengraftment in immunodeficient mice [29] and ultra-deep-sequencing of clear cell renal cell carcinoma [30]. Both datasets were previously considered for analyses by the two cancer phylogeny reconstruction methods LICHeE [31] and MIPUP [32]. Data from [29] was also used in [14] for evaluation. An interesting feature of the data in [30] is that most samples in the study present poly-occurring labels, suggesting recurrent mutations at different evolutionary stages. We recall that DISC and CASet compute dissimilarity scores, that we convert into a similarity measure subtracting their value from 1. All the analyzed trees are available in the Supplementary Materials.

To evaluate the effectiveness of the measures in real scenarios, we selected the manually curated trees, published in the corresponding original sequencing studies, for case SA501 from [29] and for patient RMH002 from [30]. We then computed similarities between these reference trees and the ones inferred by LICHeE and MIPUP, as reported in [32].

The reference RMH002 is very similar to the evolutions inferred by LICHeE and MIPUP, thus most of the measures agree on a high similarity score, as reported in Figure 8 (*Left*), with the exception of CASet_*⋃*_. The scores computed by `MP3` are higher than the others, possibly because it is the only method to correctly identify and process poly-occurring labels in the reference trees, due to the DISCovered recurring mutations. Differently from the previous analysis, the measures disagree considerably for SA501, as depicted in in Figure 8 (*Center*). Indeed, `MP3` reports a similarity value close to 0, suggesting that the considered trees are quite different, whereas the other measures report a higher similarity, especially DISC scoring up to 60% similarity.

**Figure 8:**
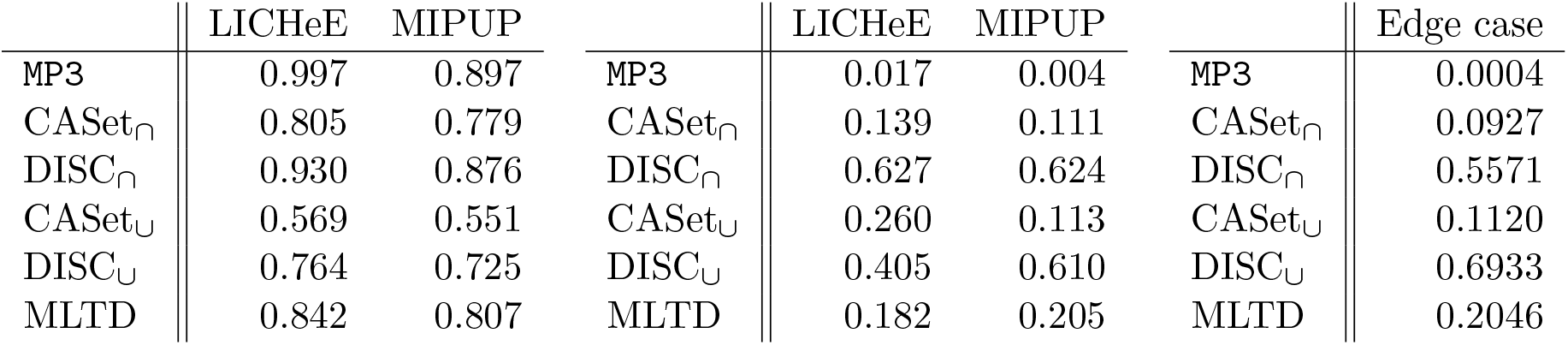
(*Left*) Similarities between the manually curated tree reported in [30] for patient RMH002 and the trees inferred by LICHeE and MIPUP. (*Center*) Similarities between the manually curated tree reported in [29] for sample SA501 and the trees inferred by LICHeE and MIPUP. (*Right*) Similarities between the manually curated tree reported in [29] for sample SA501 and the edge case with all mutations appearing in a single node.

To thoroughly investigate this behavior, we defined some naïve approaches used as a proxy to analyze some basic aspects of the trees, such as the count of pairs of labels appearing in the same node in both trees. Even with such a naïve measure, the reference tree for SA501 from [29] and the trees inferred by MIPUP and LICHeE disagree considerably. The base tree contains only 50 labels, whereas the trees inferred by LICHeE and MIPUP contain 95 and 158 labels, respectively; of these, the reference shares a total of 24 label with LICHeE and 37 with MIPUP. Most importantly, only 54 out of 1759 pairs of labels appear in the same node both in the reference and LICHeE and 124 out of 8424 in MIPUP. Such evaluations, albeit very simplistic, suggest that the trees are indeed dissimilar and thus a lower score, as provided by `MP3`, is more reasonable than a high value of similarity.

To better understand this phenomenon, we created the edge case of a single-node tree with all the 158 labels from MIPUP, and compared it against the reference SA501. The resulting values in Figure 8 (*Right*) show a high similarity score for DISC with values up to 69%, with CASet and MLTD being less influenced by this aspect with scores up to 11% and 20%. On the other hand, `MP3` clearly defines the trees as extremely dissimilar, with a score of 0.4%. Such results for trees that are clearly extremely different show a strong bias for DISC towards high similarity values.

## Discussion

We identified two major limitations in the existing methods to compare tumor phylogenies: first, they are not sensitive enough to detect even major differences in the topology of the trees, as we demonstrated with ad-hoc experiments. Second, they are not able to meaningfully compare trees where the same label is assigned to more than one node.

We addressed the latter by representing tumor phylogenies as multi-labeled trees with poly-occurring labels. Such model is best suited to cancer progression than the ones previously adopted, as it allows the same mutation to appear in multiple evolutionary events, a circumstance often occurring in real applications. Being inspired by the triplet distance for classical phylogenies, our new similarity measure correctly detects differences in the topology of the trees.

Our experiments show that our method performs very well both on synthetic and real data and, unlike the other existing tools, it is able to detect differences regarding poly-occurring labels and it suitably distinguish trees with different topologies. Moreover, when applied to hierarchical clustering, it outperforms every other method.

