## Supplementary material for "Triplet-based similarity score for fully multi-labeled trees with poly-occurring labels": Supplemetary material

#### 1 Proof of Lemma 2

Besides the five possible configurations already listed in the proof of Lemma 1, the multi-labeled model admits four additional configurations, due to the extension of the definition of LCA of two labels. In the first three additional cases, two labels are assigned to the same node and the third one to a different one. Withouth loss of generality, suppose that $a$ and $b$ label the same node. These cases are: (i) $\texttt{LCA}(a,b) = \texttt{LCA}(b,c) = \texttt{LCA}(a,c)$ and the LCA is a node in $\{a,b,c\}$, (ii) $\texttt{LCA}(a,b) \neq \texttt{LCA}(b,c) = \texttt{LCA}(a,c)$ and both LCA are nodes in $\{a,b,c\}$, and (iii) $\texttt{LCA}(a,b) \neq \texttt{LCA}(b,c) = \texttt{LCA}(a,c)$ and only $\texttt{LCA}(a,b)$ is a node in $\{a,b,c\}$. The remaining case (iv) is the simplest one in which $a$, $b$, $c$ label the same node of $T$, implying that $\texttt{LCA}(a,b) = \texttt{LCA}(b,c) = \texttt{LCA}(a,c)$.

Figure **??** reports cases (i) to (iv) from left to right.

#### 2 Experimental comparison of MP3

We show here an experimental comparison of the different versions of MP3, demonstrating the reason we decided to use MP3 $_\sigma$ as our default measure. We show in Figure S1 ($a$) that MP3 $_\sigma$ combines the best aspect of MP3 $_\cap$ and MP3 $_\cup$ while MP3 $_G$ is, as expected, the average of the two. The same result can be seen in Figure S1 ($b$), while Figure S1 ($c$) display the effect of the sigmoid, where as the trees become less similar the value move towards the union.

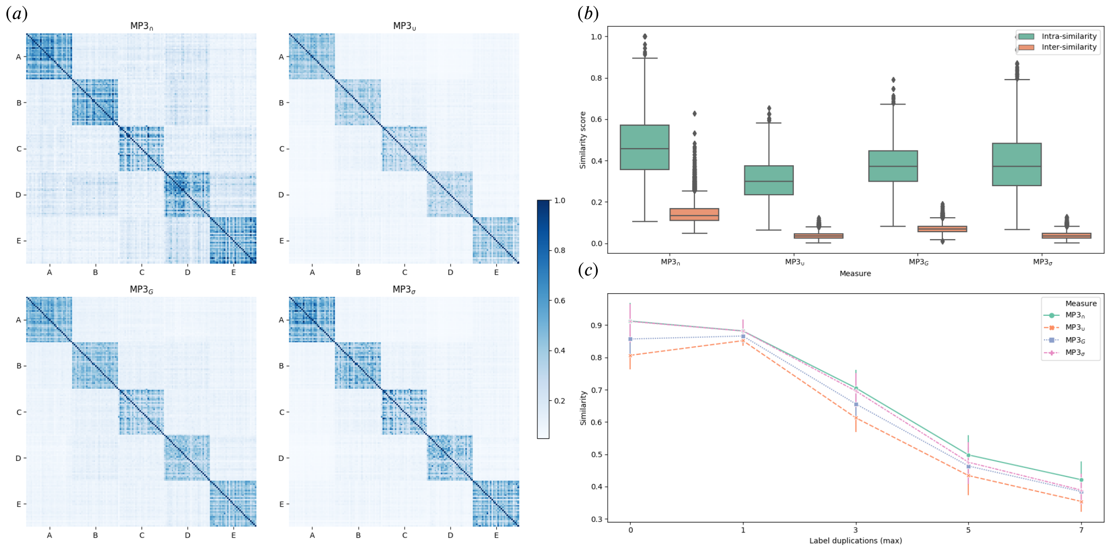


Figure S1: $(a)$ Heatmaps displaying the scores between all the 150 simulate trees from the second experimental setting. $(b)$ Distribution of the similarities between the trees in the same class (Intra-similarity) and in different classes (Inter-similarity) for the 5 classes. $(c)$ Effect of label duplication on the similarity scores. Similarities are the average of 15 trees generate from the same base with the specified value of maximum duplication from the previous experiment.

### 3 Base tree for poly-occurring label experiment

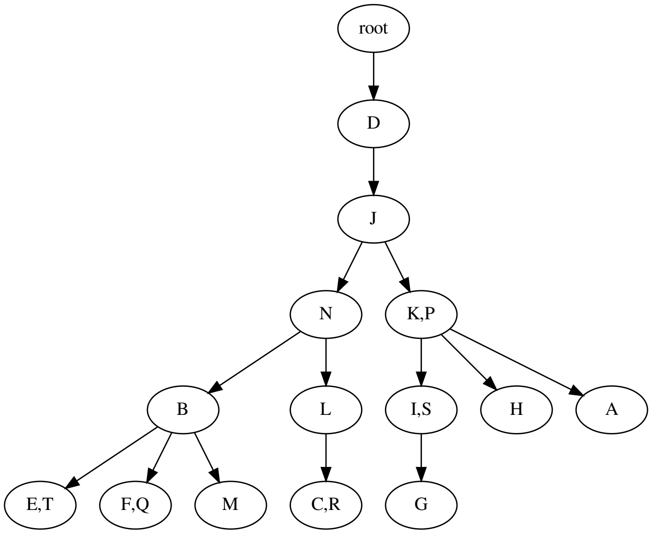


Figure S2: Base tree used for evaluating the effect of poly-occurring labels on the similarity scores.

### 4 Base trees for Exp1 and Exp2

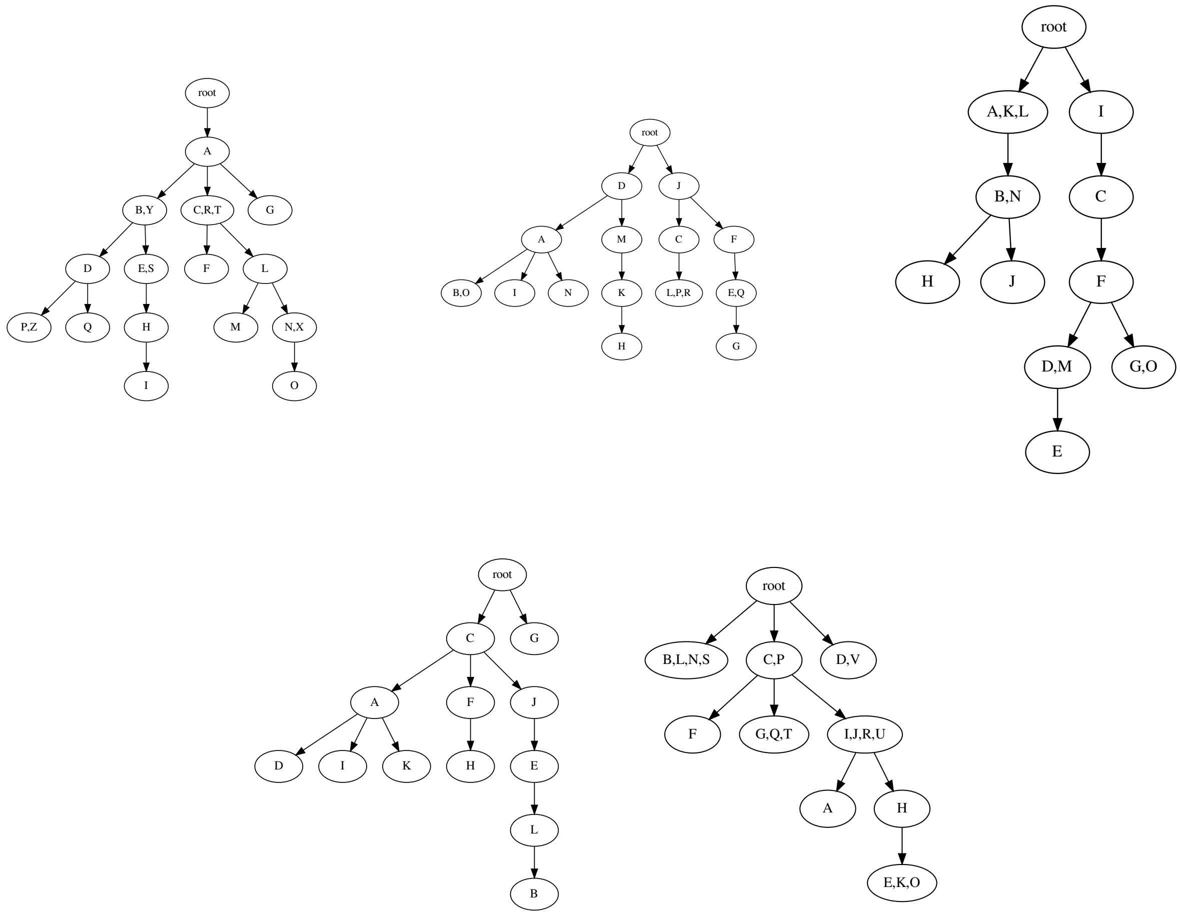


Figure S3: Base trees used in Experiments 1 and 2.

### 5 Base trees for Clustering experiment

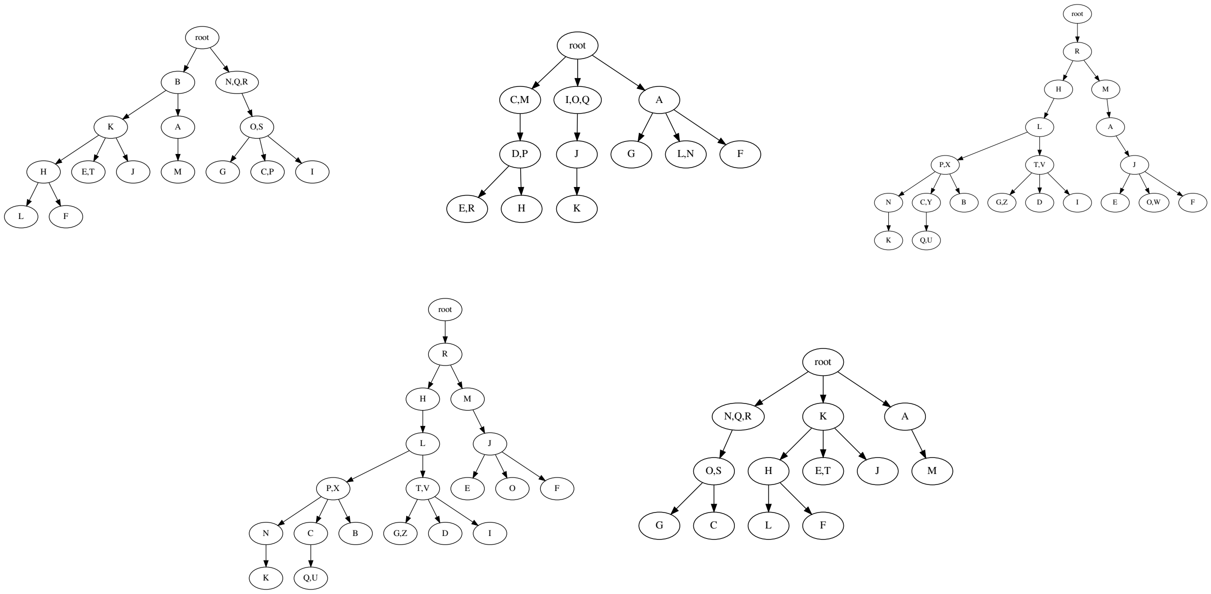


Figure S4: Base trees used in the clustering experiment. Tree 4 is a perturbation of Tree 3 and Tree 5 is a perturbation of Tree 1.

### 6 Trees for real data experiment

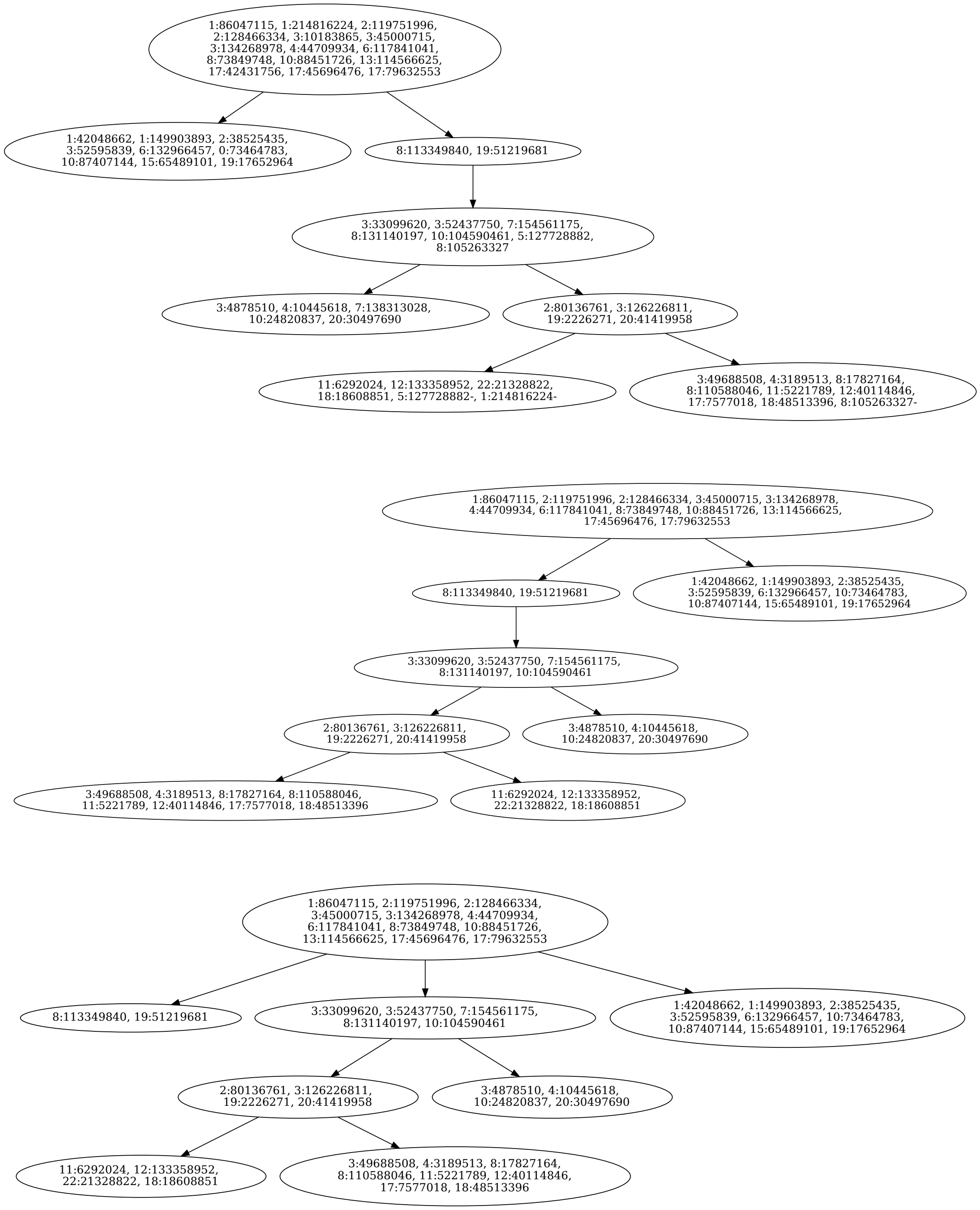


Figure S5: Trees used in the experiment on real data from Gerlinger et al. (2014). The upper tree is the base tree proposed in Gerlinger et al. (2014) for patient RMH002, the second one is the tree inferred by LICHeE, and the last one is the tree inferred by MIPUP (ipd parameter). We crafted the first tree starting from the supplementary material of the corresponding paper whereas we built the other two considering the trees reported in the MIPUP github repository.

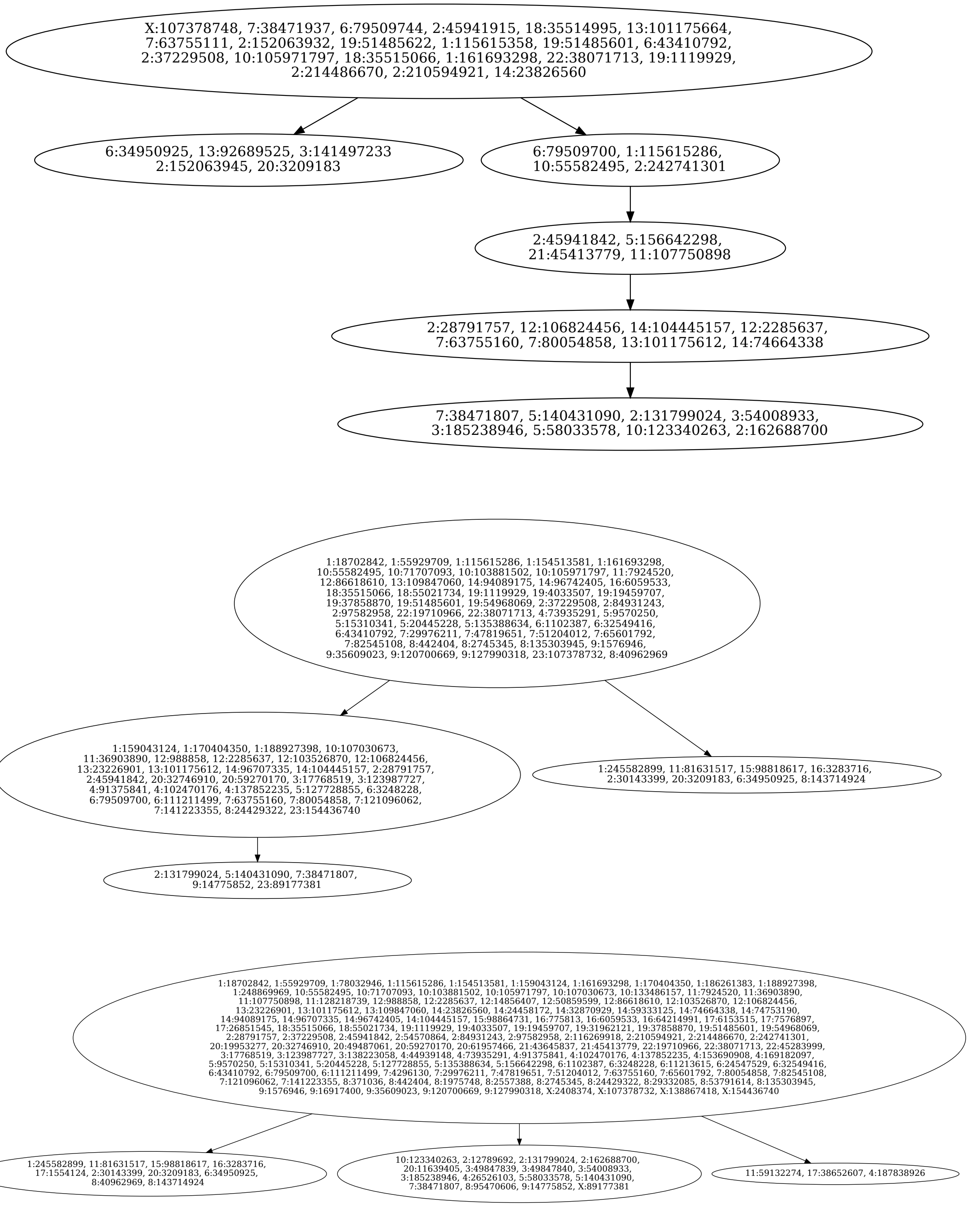


Figure S6: Trees used in the experiment on real data from Eirew et al. (2015). The upper tree is the base tree proposed in Eirew et al. (2015) for case SA501, the second one is the tree inferred by LICHeE, and the last one is the tree inferred by MIPUP. We obtained these trees from the supplementary material of DiNardo et al. (2019).

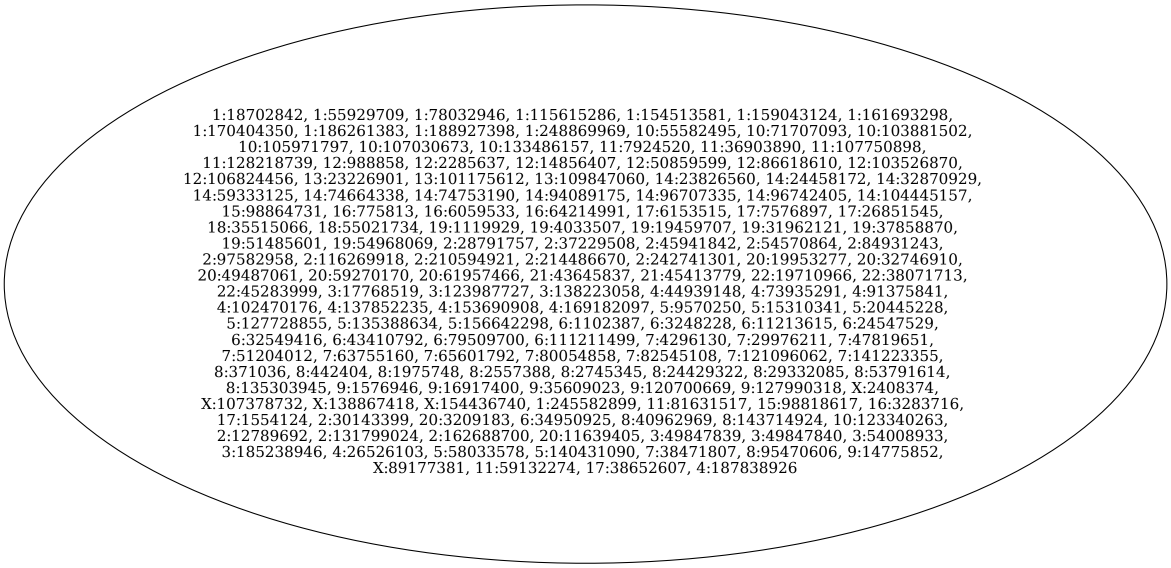


Figure S7: Edge case tree used in the experiment on real data from Eirew et al. (2015). We obtained such a tree by collapsing all nodes of the MIPUP's tree from Figure S6 in a single node.
